# IRONMAN interacts with OsHRZ1 and OsHRZ2 to maintain Fe homeostasis

**DOI:** 10.1101/2022.03.11.483574

**Authors:** Feng Peng, Chenyang Li, Chengkai Lu, Yang Li, Peng Xu, Gang Liang

## Abstract

IRONMAN is a family of small peptides which positively regulate the Fe deficiency response. However, the molecular mechanism by which OsIMA1 and OsIMA2 regulate Fe homeostasis was unclear. Here, we reveal that OsIMA1 and OsIMA2 interact with the potential Fe sensors, OsHRZ1 and OsHRZ2. OsIMA1 and OsIMA2 contain a conserved 17-amino acid C-terminal region which is responsible for the interactions with OsHRZ1 and OsHRZ2. The *OsIMA1* overexpressing plants have the increased seed Fe concentration and the reduced fertility, as observed in the *hrz1-2* loss-of-function mutant plants. Moreover, the expression trends of Fe deficiency inducible genes in the *OsIMA1* overexpressing plants are the same to those in the *hrz1-2*. Co-expression assays suggest that OsHRZ1 and OsHRZ2 promote the degradation of OsIMA1 proteins. As the interaction partners of OsHRZ1, the OsPRI proteins also interact with OsHRZ2. The conserved C-terminal region of four OsPRIs contributes to the interactions with OsHRZ1 and OsHRZ2. An artificial IMA (aIMA) derived from the C-terminal of OsPRI1 can be also degraded by OsHRZ1. Moreover, the *aIMA* overexpressing rice plants accumulate more Fe without reduction of fertility. This work establishes the link between OsIMAs and OsHRZs, and develops a new strategy for Fe fortification in rice.

## Introduction

Iron (Fe) is one of the essential micronutrients for all living organisms since Fe can mediate the redox reactions in the electron transport chain through the conversion between ferrous and ferric, which takes part in various cellular biochemical processes (Hänsch and Mendel, 2009). Fe deficiency is one of the most prevalent nutritional disorders worldwide (Mayer *et al.*, 2008). Since humans obtain Fe mainly from plants, the production of Fe-rich crops will profit human health. Fe deficiency often results in interveinal chlorosis of leaves, growth retardation and reduced crop yields (Briat *et al.*, 2015). Despite Fe is abundant in the earth’s crust, bioavailability of Fe is low as it is mainly present in the forms of insoluble hydroxides and oxides, especially in calcareous soils. One third of world’s cultivated lands are calcareous soils. Thus, Fe deficiency has become one of the factors limiting plant quality and productivity around the world.

To cope with Fe deficiency stress, plants have evolved two distinct strategies for efficient Fe uptake, the reduction strategy (strategy I) and the chelation strategy (strategy II) (Römheld and Marschner, 1986). Generally, the strategy I, which is mainly utilized in non-graminaceous plants, involves the acidification of the rhizosphere to release Fe, the reduction of Fe (III) to Fe (II) and the transport of Fe (II) (Marschner, 1995; Eide *et al.*, 1996; Robinson *et al.*, 1999). The strategy II, which is employed by graminaceous plants, is mediated by the synthesis and secretion of Fe (III) chelators, the mugineic acid (MA) family, and the translocation of MA-Fe (III) into roots. Rice (*Oryza sativa*) is a specific graminaceous species, which preferentially grows in the waterlogged field. Rice not only possesses the strategy II-based Fe-uptake system which includes the MA synthesis associated enzymes (*e. g*. OsNAS1, OsNAS2, OsNAAT1, OsDMAS1, *etc*.), the MA excretion protein (*e. g*. OsTOM1) and the MA-Fe (III) transporter (*e. g*. OsYSL15), but also partial strategy I Fe uptake system which involves the Fe (II) transporter OsIRT1 and OsIRT2 (Ishimaru *et al.*, 2006; Cheng *et al.*, 2007).

Fe deficiency directly restricts the growth and development of plants, while excessive iron causes the formation of cytotoxic reactive oxygen species (ROS) and damages cellular constituents (Brumbarova *et al.*, 2005). To maintain Fe homeostasis, plants have developed the sophisticated signaling network to regulate Fe uptake and transport. The basic helix-loop-helix (bHLH) family plays a key role in the maintenance of Fe homeostasis in rice (Kobayashi, 2019). OsIRO2 (Iron-Related bHLH Transcription Factor 2, OsbHLH56) functions as a crucial regulator of Fe homeostasis (Ogo *et al.*, 2007), which is in charge of the expression of strategy II associated genes *OsNAS1*, *OsNAS2*, *OsNAAT1*, *OsTOM1* and *OsYSL15* (Ogo *et al.*, 2011; Liang *et al.*, 2020). OsFIT (FER LIKE FE DEFICIENCY INDUCED TRANSCRIPTION FACTOR)/OsbHLH156 is an interaction partner of OsIRO2. Unlike OsIRO2 which is preferentially localized in the cytoplasm, OsFIT mainly localizes in the nucleus. In the presence of OsFIT, OsIRO2 moves to the nucleus where OsFIT and OsIRO2 form a transcription complex to activate the expression of Fe-uptake gene (Wang *et al.*, 2019; Liang *et al.*, 2020). The transcription of *OsFIT* and *OsIRO2* increases under Fe deficient conditions, but decreases under Fe sufficient conditions. OsIRO3 is a negative regulator of Fe homeostasis (Zheng *et al.*, 2010; Wang et al., 2020a; Wang et al., 2020b), and the loss-of-function of *OsIRO3* causes the up-regulation of *OsFIT* and *OsIRO2* (Li *et al.*, 2022). In contrast, OsPRI1 (Positive Regulator of Iron Homeostasis 1)/OsbHLH060, OsPRI2/bHLH058, and OsPRI3/OsbHLH059 positively regulate the expression of *OsIRO2* and *OsFIT* (Zhang *et al.*, 2017, 2020; Kobayashi *et al.*, 2019). OsHRZ1 (Haemerythrin Motif-Containing Really Interesting New Gene (RING) and Zinc-Finger Protein 1) and OsHRZ2 have been identified as potential Fe sensors playing a negative role in Fe homeostasis, which contain several hemerythrin domains for Fe binding and a RING domain for E3 ligase activity (Kobayashi *et al.*, 2013). OsHRZ1 interacts with OsPRI1, OsPRI2, and OsPRI3 and mediates their degradation via the 26S proteasome pathway (Zhang *et al.*, 2017, 2020).

IMA (IRONMAN), a family of small peptides, has been recently reported to play a positive role in the Fe deficiency response in *Arabidopsis* and rice (Hirayama *et al.*, 2018; Grillet *et al.*, 2018; Kobayashi *et al.*, 2021; Li *et al.*, 2021). Two OsIMA genes were identified in rice (Grillet *et al.*, 2018; Kobayashi *et al.*, 2021). The expression of both genes is positively regulated by OsPRI2 and OsPRI3, and negatively regulated by OsIRO3 (Wang *et al.*, 2020). However, it was still unclear how OsIMA1 and OsIMA1 activate the Fe deficiency response in rice. In the present study, we show that OsIMA1 and OsIMA2 physically interact with OsHRZ1 and OsHRZ2, and their protein stability is under the control of OsHRZs. Correspondingly, the *OsIMA1* overexpression causes the phenotypes similar to those of *hrz1-2* mutant plants. Furthermore, the overexpression of an artificial IMA derived from OsPRI1 promotes Fe accumulation in seeds, but does not lower fertility.

## Results

### OsIMA1 and OsIMA2 interact with OsHRZ1 and OsHRZ2

OsIMA1 and OsIMA2 play positive roles in Fe homeostasis since their overexpression promotes the expression of Fe deficiency inducible genes (Kobayashi *et al.*, 2021). It was reported that Arabidopsis IMAs physically interact with the C-terminal region of BTS (Li *et al.*, 2021) which is an ortholog of OsHRZ1 (Kobayashi *et al.*, 2013). OsHRZ2 is a paralog of OsHRZ1 in rice. To verify whether OsIMA1 and OsIMA2 interact with OsHRZ1 and OsHRZ2, we carried out yeast-two-hybrid assays. OsIMA1 and OsIMA2 were fused with the GAL4 DNA binding domain (BD) in the pGBKT7 vector as the baits, and the C-terminal regions of OsHRZ1 and OsHRZ2 with the GAL4 activation domain (AD) in the pGADT7 vector as the preys. As shown in the growth of yeast, both OsIMA1 and OsIMA2 interact with the C-terminal regions of OsHRZ1 and OsHRZ2 (Figure 1A).

**Figure 1.**
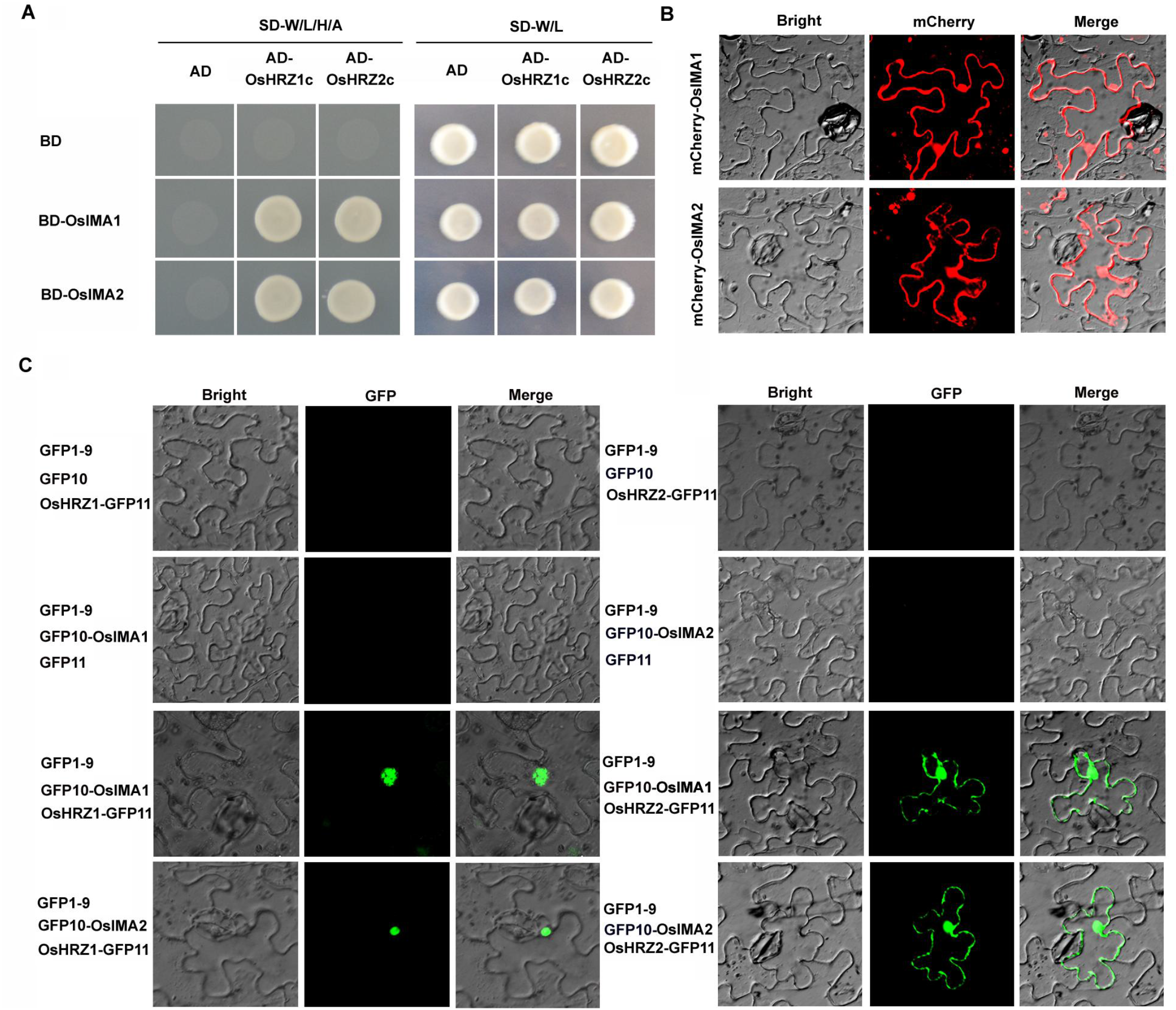
OsIMA1 and OsIMA2 interact with OsHRZ1 and OsHRZ2. (A) OsIMAs interact with the C-terminal regions of OsHRZs in yeast. The full-length OsIMAs were fused with BD, and the C-terminal regions of OsHRZs with AD. Yeast co-transformed with different BD and AD plasmid combinations was spotted. Growth on selective plates lacking leucine, tryptophan, adenine, and histidine (−4) or lacking leucine and tryptophan (−2) is shown. (B) Subcellular localization of OsIMA1 and OsIMA2. mCherry was fused with the N-end of OsIMAs. Transient expression assays were performed in *N. benthamiana* leaves. (C) Interaction of OsIMAs and OsHRZs in plant cells. Tripartite split-sfGFP complementation assays were performed. OsIMAs were fused with GFP10, and OsHRZs with GFP11. The combinations indicated were introduced into Agrobacterium and co-expressed in *N. benthamiana* leaves.

OsHRZ1 protein localizes in the nucleus, and OsHRZ2 in both the nucleus and cytoplasm (Kobayashi *et al.*, 2013). To investigate the subcellular localization of OsIMA1 and OsIMA2, mCherry was tagged to the N-end of OsIMA1 and OsIMA2 respectively and expressed in tobacco leaves. As shown in Figure 1B, both OsIMA1 and OsIMA2 were present in the nucleus and cytoplasm. To further confirm the location where the protein interactions occur, we employed the tripartite split-GFP system monitoring the localization of protein complex. The GFP10 fragment was fused with OsIMA proteins in their N-end (GFP10-OsIMAs) and the GFP11 with OsHRZs in their C-end (OsHRZs-GFP11). When GFP10-OsIMA1/2 and OsHRZ1-GFP11 were transiently co-expressed with GFP1-9 in tobacco leaves, the GFP signal was only visible in the nucleus of transformed cells (Figure 1C). By contrast, when GFP10-OsIMA1/2 and OsHRZ2-GFP11 were transiently co-expressed with GFP1-9, the GFP signal was visible in both the nucleus and cytoplasm. Taken together, our data suggest that OsIMAs physically interact with OsHRZs in plant cells.

### The C-terminal region of OsIMAs accounts for the interactions with OsHRZ1 and OsHRZ2

The IMAs feature a conserved C-terminal region (Figure 2A; Grillet *et al.*, 2018). We wanted to know whether the C-terminal region is responsible for their interactions with OsHRZs. We performed yeast-two-hybrid assays. OsIMA peptides were divided into two parts, the N-terminal region and the C-terminal 17-amino acid region, and respectively fused to the BD in the pGBK-T7 vector as baits. Yeast growth assays indicated that their C-terminal regions, but not the N-terminal regions, interact with OsHRZs (Figure 2B).

**Figure 2.**
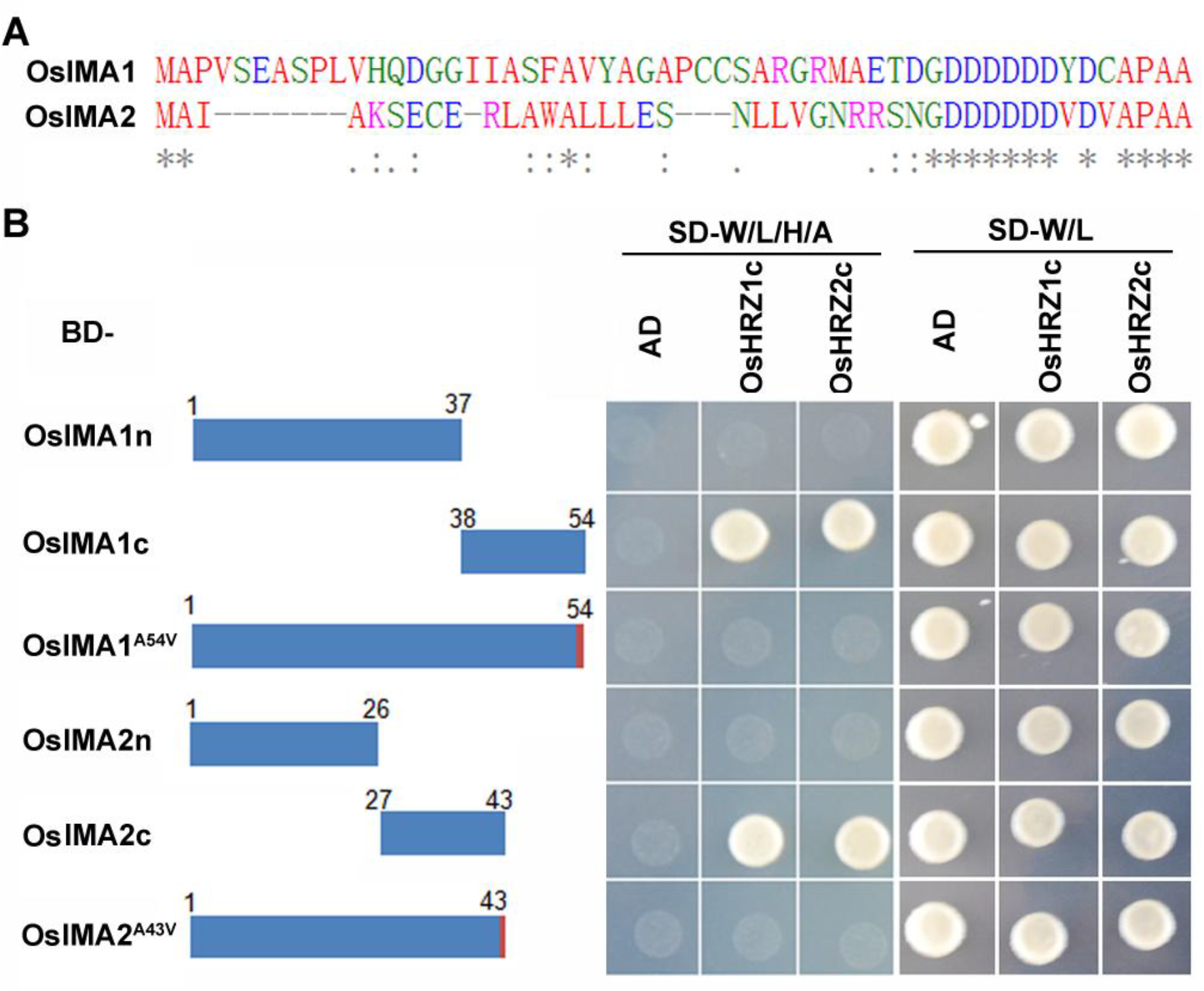
The last amino acid of OsIMA1 and OsIMA2 is crucial for interactions with OsHRZ1 and OsHRZ2. (A) Alignment of amino acid sequences of OsIMAs. The full-length amino acid sequences of OsIMAs were aligned with Clustal Omega online (https://www.ebi.ac.uk/Tools/msa/clustalo/). (B) Yeast-two-hybrid assays. The truncated or mutated OsIMAs were fused with BD, and the C-terminal regions of OsHRZs with AD. Yeast co-transformed with different BD and AD plasmid combinations was spotted. Growth on selective plates lacking leucine, tryptophan, adenine, and histidine (−4) or lacking leucine and tryptophan (−2) is shown.

The last amino acid A of Arabidopsis IMAs is crucial for their interactions with BTS (Li *et al.*, 2021). The last amino acid of OsIMAs is also A. We asked whether the same case occurs in rice. We generated the full-length OsIMAs with their last amino acid changed from A to V, and fused them with the BD as baits. Yeast growth indicated that the mutation of last amino acid A disrupted the interactions between OsIMAs and OsHRZs (Figure 2B). These data suggest that the C-terminal regions of OsIMAs contribute to their interactions with OsHRZs, and the last amino acid A is necessary.

### *OsIMA1* overexpressing plants mimic the *hrz1-2* mutant plants

To further investigate how OsIMAs regulate the Fe deficiency response, we generated *OsIMA1* overexpressing transgenic plants in which the *OsIMA1* gene was driven by the maize ubiquitin promoter (Figure S1). The *OsIMA1* transgenic plants displayed the reduced fertility, which is also observed in the *hrz1-2* loss-of-function mutant plants (Figure 3A). Measurement of Fe concentration indicated that the *OsIMA1* overexpressing plants accumulated much more Fe in the seeds than the wild type plants, as did the *hrz1-2* mutant plants (Figure 3B). Considering that the *OsIMA1* overexpressing plants phenocopied the *hrz1-1* mutant plants, we then compared the expression of several Fe deficiency inducible genes. In the Fe deficiency response signaling pathway, *OsIRO3* and *OsIRO2* are two major transcription factors which are considerably up-regulated under Fe deficient conditions. Additionally, the strategy II associated genes, such *OsNAS1*, *OsNAS2*, *OsTOM1* and *OsYSL15*, are strongly induced by Fe deficiency. Rice plants were grown in Fe sufficient solution for two weeks, and then their roots were separated and used for RNA extraction. Examination of transcript abundance indicated that the expression of those Fe deficiency inducible genes was up-regulated significantly in both the *OsIMA1* overexpressing plants and *hrz1-2* mutant plants compared with the wild type plants (Figure 3C).

**Figure 3.**
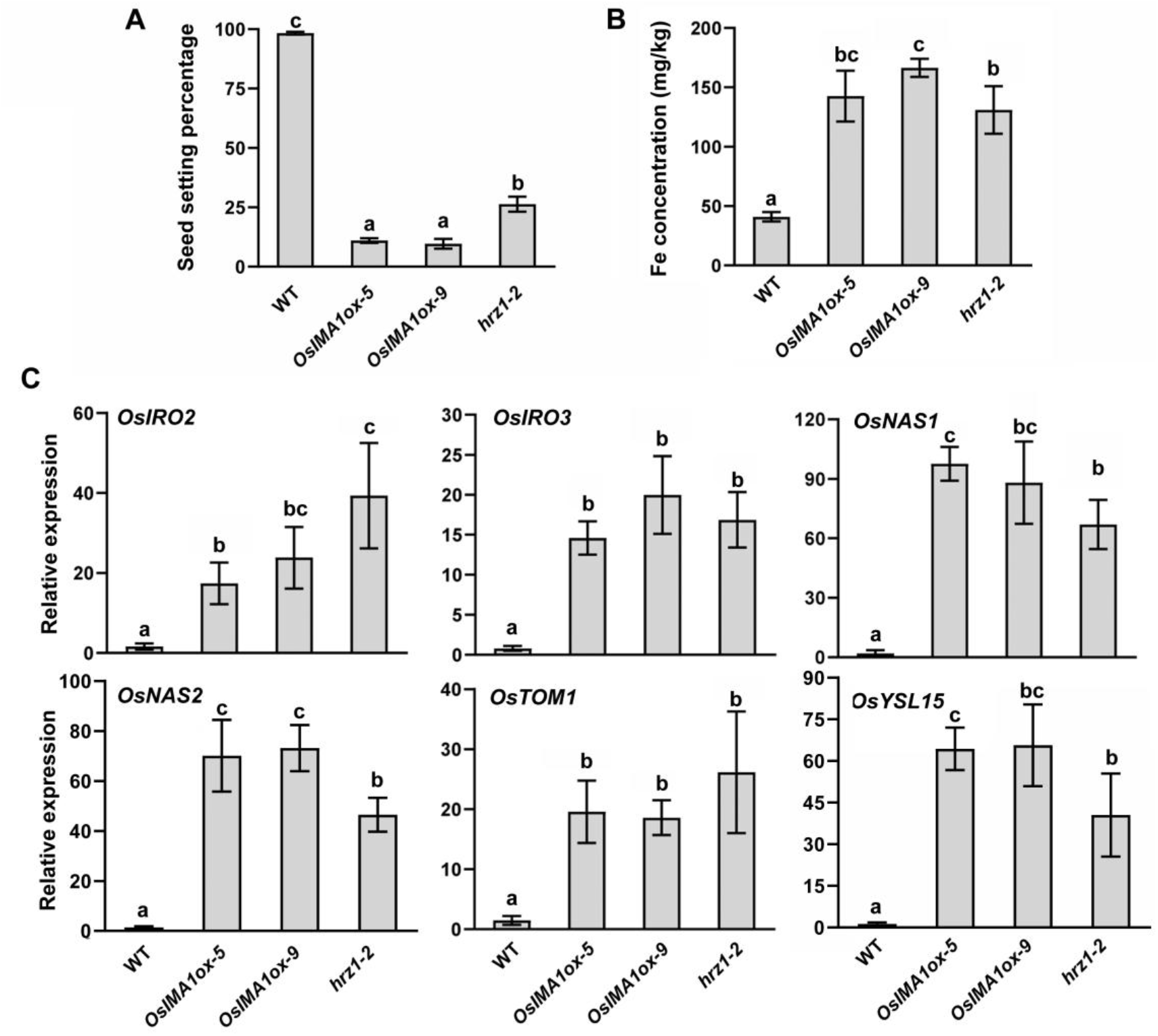
*OsIMA1* overexpressing plants mimic the *hrz1-2* loss-of-function mutant. (A) Seed setting percentage. Data represent means ± standard deviation (SD) (*n* = 3). Different letters above each bar indicate statistically significant differences (ANOVA, P < 0.05). (B) Fe concentration. Brown seeds were used for Fe measurement. Data represent means ± standard deviation (SD) (*n* = 3). Different letters above each bar indicate statistically significant differences (ANOVA, P < 0.05). (C) Expression of Fe deficiency inducible genes. Rice plants were grown in Fe sufficient solution for two weeks, and roots were used for RNA extraction and qRT-PCR. Data represent means ± standard deviation (SD) (*n* = 3). Different letters above each bar indicate statistically significant differences (ANOVA, P < 0.05).

### OsHRZ1 and OsHRZ2 promote the degradation of OsIMAs

OsHRZ1 and OsHRZ2 have a RING domain which possesses E3 ligase activity and several proteins have been reported to be degraded by OsHRZ1 and OsHRZ2 (Zhang *et al.*, 2017, 2020; Guo *et al.*, 2022). To investigate whether OsHRZ1 and OsHRZ2 also facilitate the degradation of OsIMAs, we used OsIMA1 as a representative and carried out the transient expression assays in tobacco leaves. The C-end of OsHRZ1 and OsHRZ2 was fused with the GFP tag. mCherry-OsIMA1 was co-expressed with GFP, OsHRZ1-GFP, and OsHRZ2-GFP respectively in tobacco leaves. Immunoblot analysis indicated that the protein levels of mCherry-OsIMA1 were significantly lower in the presence of OsHRZ1-GFP or OsHRZ2-GFP compared with in the presence of GFP (Figure 4). These data suggest that OsHRZ1 and OsHRZ2 accelerate the degradation of OsIMA1 and OsIMA2.

**Figure 4.**
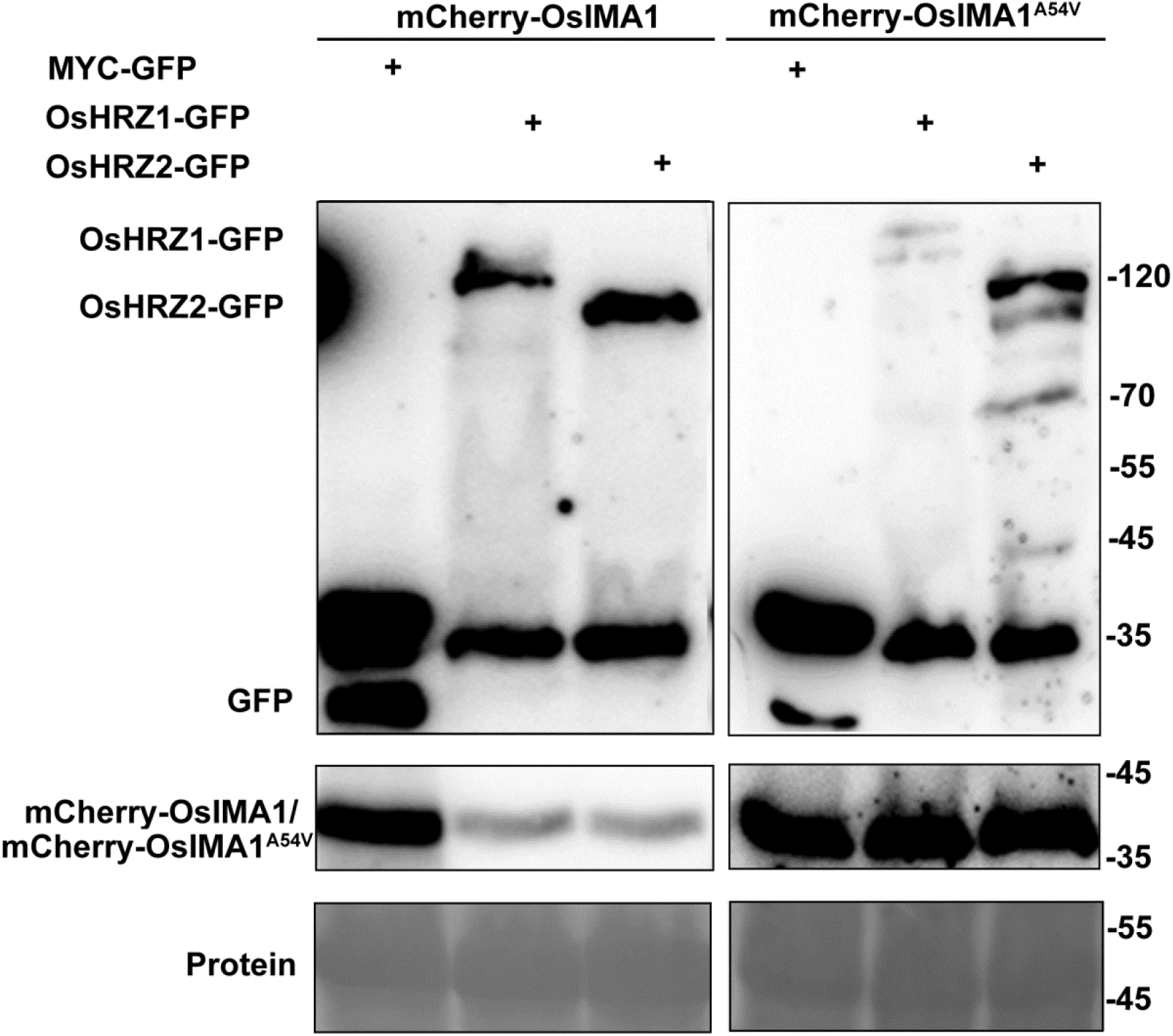
Both OsHRZ1 and OsHRZ2 promote the degradation of OsIMA1. mCherry-OsIMA1 or mCheryy-OsIMA1 was coexpressed with MYC-GFP, OsHRZ1-GFP, and OsHRZ2-GFP, respectively. Total protein was extracted and immunblotted with anti-GFP antibody or anti-mCherry antibody. Ponceau staining shows equal loading. Protein molecular weight (in kD) is indicated.

Given that the last amino acid of OsIMA1 and OsIMA2 is responsible for their interactions with OsHRZ1 and OsHRZ2, we speculated that the protein stability of OsIMA1^A54V^ would not be affected by OsHRZ1 and OsHRZ2. The mCherry tag was linked with the N-end of OsIMA1^A54V^, and then used for co-expression assays. As expected, OsHRZ1 and OsHRZ2 could not degrade OsIMA1^A54V^ (Figure 4). These data suggest that OsHRZ1 and OsHRZ2 degrade OsIMAs in a protein interaction dependent manner.

### The C-terminal region of OsPRI1 interacts with OsHRZ1 and OsHRZ2

The rice bHLH IVc subgroup consists of four members, and three of them (OsPRI1, OsPRI2, and OsPRI3) interact with OsHRZ1 (Zhang *et al.*, 2017, 2020). To verify if the fourth member OsPRI4 also interacts with OsHRZ1, we tested their interaction using the yeast-two-hybrid system. Yeast growth indicated that OsPRI4 and OsHRZ1 interact with each other. Due to OsHRZ2 is a paralog of OsHRZ1, we wanted to know if OsHRZ2 is also an interaction partner of these four OsPRI proteins. Protein interaction tests indicated that these four OsPRI proteins also interact with OsHRZ2 (Figure 5A).

**Figure 5.**
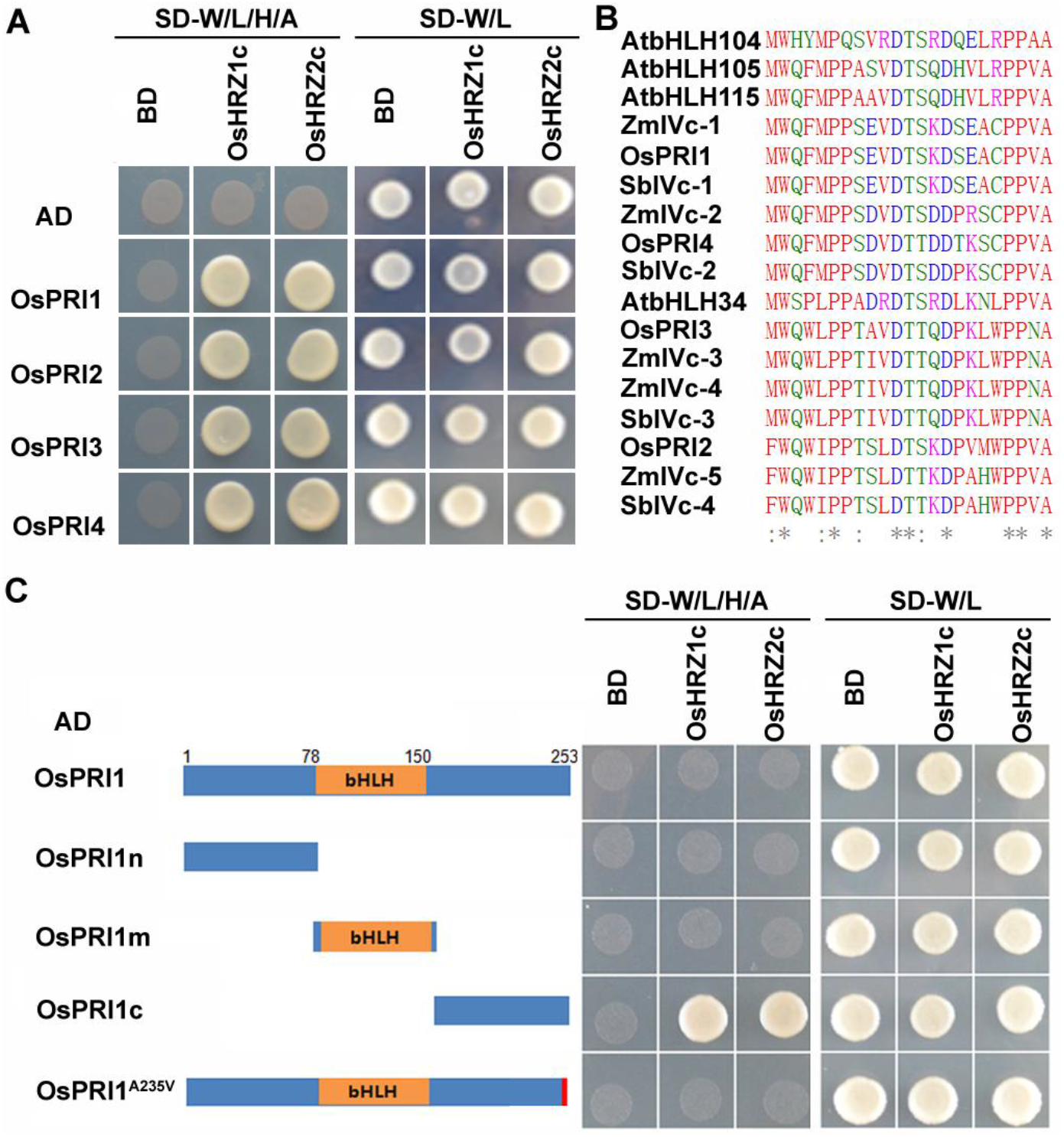
The C-terminal region of OsPRIs interacts with OsHRZs. (A) All four OsPRI proteins interact with both OsHRZ1 and OsHRZ2. The full-length OsPRIs were fused with AD, and the C-terminal regions of OsHRZs with BD. Yeast co-transformed with different BD and AD plasmid combinations was spotted. Growth on selective plates lacking leucine, tryptophan, adenine, and histidine (−4) or lacking leucine and tryptophan (−2) is shown. (B) The C-terminal regions of bHLH IVc proteins from different plants. bHLH IVc proteins in *Arabidopsis thaliana*, *Oryza sativa*, *Zea mays*, and *Sorghum bicolor*, were used for analysis. AtbHLH34 (AT3G23210), AtbHLH104 (AT4G14410), AtbHLH105 (AT5G54680), AtbHLH115 (AT1G51070), OsPRI1 (LOC_Os08g04390), OsPRI2 (LOC_Os05g38140), OsPRI3 (LOC_Os02g02480), OsPRI4 (LOC_Os07g35870), ZmIVc-1 (ZmPHJ40.04G070000), ZmIVc-2 (ZmPHJ40.07G193300), ZmIVc-3 (ZmPHJ40.04G350300), ZmIVc-4 (ZmPHJ40.05G151400), ZmIVc-5 (ZmPHJ40.06G183000), SbIVc-1 (SbiSC187.07G031300), SbIVc-2(SbiSC187.02G307200), SbIVc-3(SbiSC187.04G009900), SbIVc-4 (SbiSC187.09G150000). (C) The C-terminal region of OsPRI1 interacts with OsHRZ1 and OsHRZ2. The truncated or mutated OsPRI1 were fused with AD, and the C-terminal regions of OsHRZs with BD. Yeast co-transformed with different BD and AD plasmid combinations was spotted. Growth on selective plates lacking leucine, tryptophan, adenine, and histidine (−4) or lacking leucine and tryptophan (−2) is shown.

The alignment of four OsPRI proteins shows that they share a conserved C-terminal region (Figure 5B). We then asked whether their interactions with OsHRZ1 and OsHRZ2 depend on their C-terminal regions. Considering the high similarity of their C-terminal regions of bHLH IVc proteins across different plant species (Figure 5B), OsPRI1 was chosen as a representative for further analysis. OsPRI1 was divided into three parts, the N-terminal (OsPRI1n), the bHLH domain (OsPRI1m), and the C-terminal (OsPRI1c), and fused with the AD. OsHRZ1c and OsHRZ2c were fused with the BD. Interaction tests indicated that only OsPRI1c could interact with OsHRZ1c and OsHRZ2c (Figure 5C). To further investigate whether the last amino acid A of OsPRI1 is crucial for the interactions, the A was substituted for V in the full length of OsPRI1. The results demonstrated that the substitution disrupted the interaction with OsHRZ1c and OsHRZ2c. These data suggest that the C-terminal region of OsPRIs is required for the interactions with OsHRZ1 and OsHRZ2.

### Generation of Fe-fortified rice grains by manipulating an artificial IMA derived from OsPRI1

The overexpression of *OsIMA1* caused the Fe over-accumulation in grains, but the reduced fertility. Because the reduction of fertility is a disadvantageous factor for yield, we attempted to develop an artificial IMA which would increase seed Fe concentration but not reduce fertility. Given the functional similarities between OsIMAs and the C-terminal regions of OsPRIs, the C-terminal region of OsPRI1 was designed as an artificial IMA (aIMA) (Figure S2A). Having confirmed that aIMA interacting with OsHRZ1 and OsHRZ2 (Figure 5C), we wondered whether aIMA could be degraded by OsHRZ1 and OsHRZ2. The mCherry tag was fused to the N-end of aIMA and used for transient co-expression assays in tobacco leaves. Compared with the GFP, both OsHRZ1-GFP and OsHRZ2-GFP caused the damage to mCherry-aIMA proteins, suggesting that OsHRZ1 and OsHRZ2 favor the degradation of aIMA (Figure 6A). To further investigate whether aIMA can increase Fe accumulation in seeds, we generated transgenic rice overexpressing aIMA (Figure S2B, C). We did not observe visible fertility difference between the wild type and *aIMA* overexpressing plants (Figure S2D). Then, we determined the Fe concentration of brown seeds, finding that the Fe concentration was around two-fold higher in the *aIMA* overexpressing plants than in the wild type (Figure 6B). Taken together, our data suggest that the artificial IMA strategy is effective on improving Fe concentration of rice grains.

**Figure 6.**
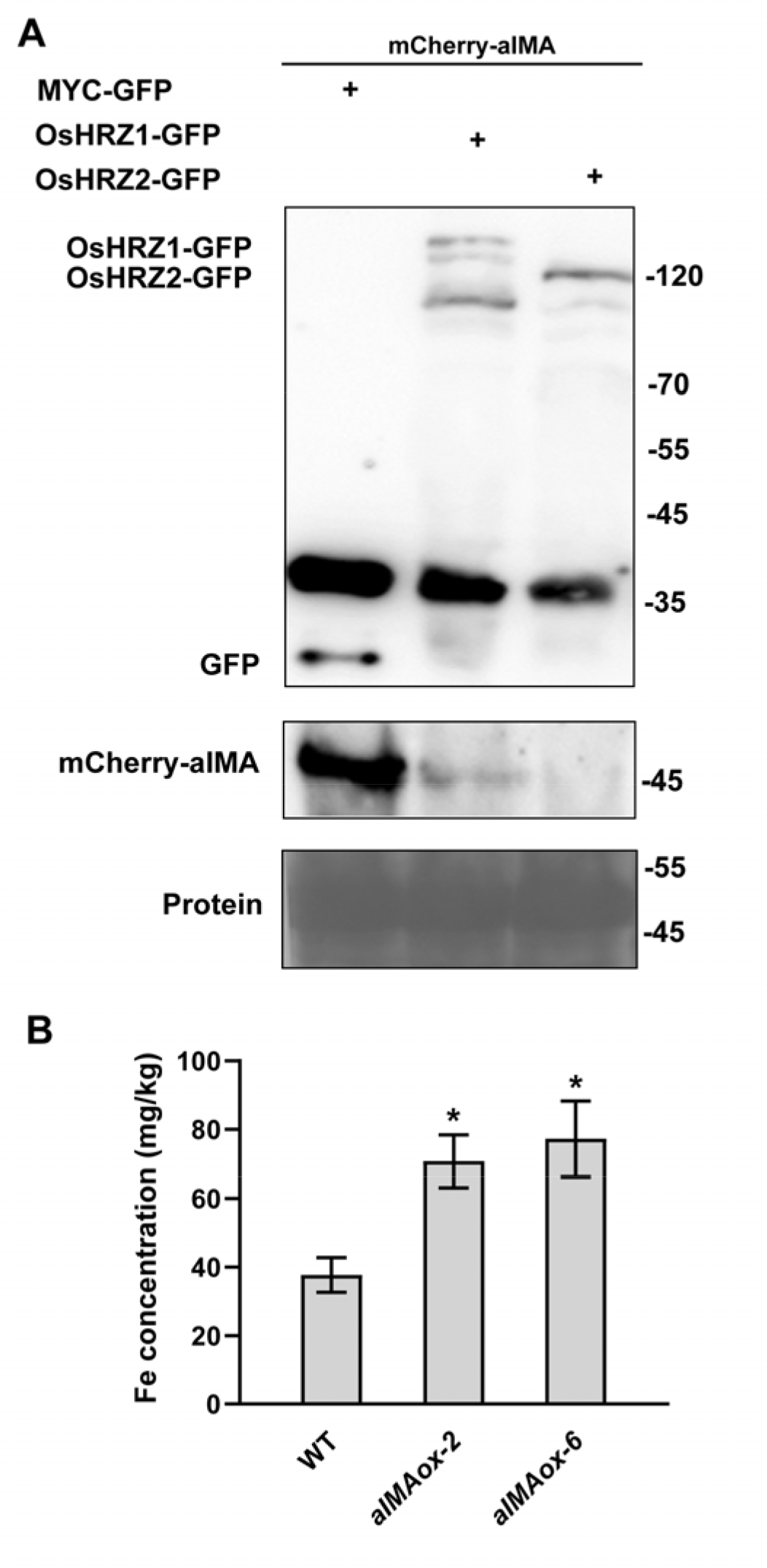
Overexpression of *aIMA* causes Fe over-accumulation in seeds. (A) Degradation of aIMA by OsHRZs. mCherry-aIMA was coexpressed with MYC-GFP, OsHRZ1-GFP, and OsHRZ2-GFP, respectively. Total protein was extracted and immunblotted with anti-GFP antibody or anti-mCherry antibody. Ponceau staining shows equal loading. Protein molecular weight (in kD) is indicated. (B) Fe concentration. Brown seeds were used for Fe measurement. Data represent means ± standard deviation (SD) (*n* = 3). The asterisk indicates a significant difference from the wild type as determined by Student’s t Test (*P* < 0.05).

## Discussion

Recently, great progress has been made in the Fe deficiency response signaling pathway in rice. Many transcription factors, especially bHLH proteins, play key roles in the Fe signaling transduction (Kobayashi, 2019). OsHRZ1 and OsHRZ2 are two potential Fe sensors which negatively regulate Fe homeostasis (Kobayashi *et al.*, 2013). IMAs are a class of small peptides which are conserved across angiosperms (Grillet *et al.*, 2018). Two OsIMAs exist in rice, and their overexpression causes the constitutive activation of Fe deficiency inducible genes (Kobayashi *et al.*, 2021). However, the underlying molecular mechanism by which OsIMA1 and OsIMA2 mediate Fe signaling remains to be clarified. Here, we suggest that OsIMAs and OsHRZs regulate antagonistically the Fe deficiency response. Based on the regulatory mechanism of OsIMAs, we developed a strategy for generation of Fe fortified rice by manipulating an artificial IMA peptides.

OsHRZ1 and OsHRZ2 contain 1-3 haemerythrin domains and one RING domain which endow them the Fe binding ability and E3 ligase activity, respectively (Kobayashi *et al.*, 2013). Here, we revealed that OsHRZ1 and OsHRZ2 interact with OsIMAs and accelerate the degradation of OsIMAs. Recently, it was reported that OsHRZ1 interacts with and mediates the degradation of three bHLH IVc proteins, OsPRI1, OsPRI2, and OsPRI3 (Zhang *et al.*, 2017, 2020). We further confirmed that all four bHLH IVc members interact with both OsHRZ1 and OsHRZ2 (Figure 5A). Given that both OsHRZ1 and OsHRZ2 have E3 ligase activity and degrade their substrates (Kobayashi *et al.*, 2013; Zhang *et al.*, 2017; Guo *et al.*, 2022), it is very likely that OsHRZ2 also mediates the degradation of bHLH IVc proteins. Although bHLH IVc proteins are the positive regulators of the Fe deficiency response, their transcription is barely induced by Fe deficiency (Zhang *et al.*, 2017, 2020; Kobayashi *et al.*, 2019). Thus, it is expected that their protein levels increase in response to Fe deficiency. It is noteworthy that the transcript abundance of *OsHRZ1* is also increased under Fe deficient conditions. In order to maintain stable levels of bHLH IVc proteins, the protein stability or activity of OsHRZ1 must be reduced under Fe deficient conditions. Although OsHRZ1 and OsHRZ2 can bind Fe ions, their protein stability seems not to respond to Fe status according to the cell free degradation assays in vitro (Kobayashi *et al.*, 2013). A latest study found that OsHRZ2 protein increases in response to Fe deficiency as shown in the *OsHRZ2-GFP* overexpressing transgenic plants (Guo *et al.*, 2022). BTS is a homolog of OsHRZ1 in Arabidopsis, and its protein stability is enhanced in the absence of Fe as shown in the protein translation in wheat germ extract (Selote *et al.*, 2015). Thus, the protein stability of OsHRZ1 and OsHRZ2 is either irrespective to Fe status or increased under Fe deficiency conditions. It was an open question how rice plants promote the accumulation of bHLH IVc proteins to activate the Fe deficiency response. In Arabidopsis, IMAs interfere with the interactions between bHLH IVc members and BTS since IMAs and bHLH IVc share a similar C-terminal region which contributes to the interactions with BTS (Li *et al.*, 2021). We indicated that the C-terminal regions of both OsIMAs and OsPRIs contain an OsHRZs interacting domain, which makes it possible that OsIMAs compete with OsPRIs for interacting with OsHRZs. This hypothesis is further supported by the fact that the *OsIMA1* overexpressing plants phenocopy the *hrz1-2* mutant plants (Figure 3). Thus, it is very likely that OsIMAs interfere with the interactions between OsPRIs and OsHRZs, hence stabilizing the OsPRI proteins and activating the Fe deficiency response.

The reciprocal regulation between OsIMAs and OsHRZs is crucial for the maintenance of Fe homeostasis. Under Fe deficient conditions, the transcription of both OsIMAs and OsHRZs is enhanced (Kobayashi *et al.*, 2013, 2021). We show that OsHRZ1 and OsHRZ2 facilitate the degradation of OsIMA1 (Figure 4). On the other hand, *OsIMA1* overexpression enhances the transcription of *OsHRZ1*, and the RNA interference of *OsHRZs* promotes the transcription of *OsIMA1* and *OsIMA2* (Kobayashi *et al.*, 2021). No matter the overexpression of *OsIMA1* or the loss-of-function of *OsHRZ1*, both cause the Fe over-accumulation and the reduced fertility in rice (Figure 3A). The disruption of balance between OsIMA1 and OsHRZ1 results in the disorder of Fe homeostasis and the Fe toxicity. The transcript abundance of *OsIMAs* is relatively low under Fe sufficient conditions, but extremely high under Fe deficient conditions (Kobayashi *et al.*, 2021). When *OsIMA1* was overexpressed under Fe sufficient conditions, rice plants accumulated excessive Fe (Figure 3B; Kobayashi *et al.*, 2021). Therefore, appropriate OsIMA levels are required for the maintenance of Fe homeostasis. In addition to degradation by OsHRZs, the transcription of *OsIMAs* is positively regulated by OsPRIs (Kobayashi *et al.*, 2021). The balance between the positive regulators (OsPRIs) and the negative regulators (OsHRZs) maintains the appropriate levels of OsIMAs.

Fe-deficiency anemia is one of the most prevalent human micronutrient deficiencies around the world. Breeding staple crops with abundant Fe is an ideal way to cope with Fe deficiency anemia. Fe fortification in rice grains has been accomplished by introducing Fe-homeostasis associated genes (Masuda *et al.*, 2013). In the *OsIMA1* overexpressing plants, the expression of Fe uptake associated genes shows marked enhancement irrespective of Fe status, which explains the increased Fe accumulation in grains when plants are grown in Fe sufficient soil. It was reported that the Fe over-accumulation is related with the embryo lethality of various *bts* mutants (Selote *et al.*, 2015). In contrast to the complete infertility of *bts* null mutants, the other weak *bts* mutants with reduced induction of *BTS* have slight embryo lethality. Although the increased Fe accumulation is advantageous for Fe fortification, the reduced fertility is disadvantageous for rice yield. The inhibitory effect of OsIMAs on OsHRZ1 provides an alternative approach to increasing Fe accumulation without reducing yield. We developed an artificial small peptide, aIMA, which possesses the ability to interact with OsHRZs and can be degraded by OsHRZs. Indeed, the increased Fe accumulation and normal fertility were achieved in the transgenic plants overexpressing aIMA peptides (Figure 6). Unlike the strong increase of Fe concentration in the *OsIMAox* plants, the moderate Fe increase was detected in the *aIMAox* plants. The major difference between OsIMAs and aMIA is that OsIMA are rich in aspartic acid. Grillet *et al.* (2018) revealed that the aspartic acid stretch contributes to the affinity of IMAs for Fe ions. Since the stability of haemerythrin domain containing proteins is affected when Fe ions are present (Salahudeen *et al.*, 2009; Vashisht *et al.*, 2009; Selote *et al.*, 2015), it is plausible that OsIMAs deliver Fe ions to OsHRZs and then result in the inactivity of OsHRZs. It might be the reason why OsIMA1 has stronger activation to Fe deficiency inducible genes than aIMA. This also raises the possibility of further improving Fe accumulation by optimizing the amino acids of aIMA. We noted that the bHLH IVc proteins share the conserved C-terminal region across different plant species (Figure 5B). Thus, the artificial IMA strategy can apply to other plant species beyond rice. Our exploration of an artificial IMA peptide provides a new strategy for Fe fortification in crops.

## Materials and methods

### Plant materials and growth conditions

Rice (*Oryza sativa* L. cv. Nipponbare) was used in this study. Rice plants were grown in Crops Conservation and Breeding Base of XTBG, in Mengla county of Yunnan province. For hydroponic culture assays, half-strength Murashige and Skoog (MS) media (pH5.6-5.8) with 0.1 mM EDTA-Fe (III). The nutrient solution was exchanged every 3 d. Plants were grown in a growth chamber at 28°C during the day and at 20°C during the night.

### qRT-PCR

Total RNA was extracted from rice roots or shoots. RNA samples were reverse transcribed using an RT Primer Mix (oligo dT) and PrimeScript RT Enzyme Mix for qPCR (TaKaRa, Japan) following the manufacturer's protocol. qRT-PCR was performed on a Light-Cycler 480 real-time PCR machine (Roche, Switzerland) by using PrimeScript™ RT reagent (Perfect Real Time) Kit (TaKaRa, Japan). All PCR amplifications were performed in triplicate, with the *OsACTIN1* and *OsOBP* as the internal controls to normalize the samples. Primer sequences used for qRT-PCR are listed in Supplemental Table S1.

### Metal concentration measurement

Seeds were dried at 65°C for one week. About 500 mg dry weight of them were digested with 5 ml of 11 M HNO_3_ for 3 h at 185°C and 2 ml of 12 M HClO_4_ for 30 min at 220°C. The concentration of Fe was measured with an Inductively Coupled Plasma Mass Spectrometry (ICP-MS, Japan). Three biological replicates were performed for analysis.

### Yeast two-hybrid assays

The yeast two-hybrid assays were carried out according to the manufacturer's protocol. The C-terminal regions of OsHRZ1 and OsHRZ2 were respectively cloned into the pGADT7 or pGBKT7 plasmids. The full-length or truncated OsIMAs were respectively cloned into pGBKT7 plasmid. The GAD-PRIs vectors were described previously (Zhang *et al.*, 2020). Vectors were transformed into yeast strain Y2HGold (Clontech, Japan). Yeast transformation was performed according to the Yeastmaker Yeast Transformation System 2 User Manual (Clontech, Japan). The primers used are listed in Supplemental Table S1.

### Generation of transgenic plants

For generation of the *OsIMA1* overexpression construct, the full-length CDS of OsIMA1 was cloned downstream of maize ubiquitin promoter in the pUN1301 binary vector. For generation of the *aIMA* overexpression construct, the C-terminal region of OsPRI1 was cloned downstream of maize ubiquitin promoter in the pUN1301 binary vector. The constructs were introduced into the *Agrobacterium tumefaciens* strain EHA105 and then used for rice transformation. The transgenic plants with hygromycin were selected and used for examination of transgene levels. T3 transgenic plants were used for analysis.

### Tripartite split-GFP complementation assays

Tripartite split-GFP complementation assays were conducted as previously reported (Liang *et al.*, 2020). Briefly, the full-length OsIMAs were respectively cloned into pTG-GFP10, and the full-length OsHRZs to pTG-GFP11. *A. tumefaciens* strain EHA105 was used in the transient expression experiments. The various combinations of agrobacterial cells were infiltrated into 3-week-old *Nicotiana benthamiana* leaves by an infiltration buffer (0.2 mM acetosyringone, 10 mM MgCl_2_, and 10 mM MES, pH 5.6). The abaxial sides of leaves were injected with 20 μM β-estradiol 24 h before observation. GFP fluorescence was photographed on an OLYMPUS confocal microscope.

### Degradation assays

OsHRZ1 and OsHRZ2 were fused with the C-end of GFP, and driven by the CaMV 35S promoter. OsIMA1, OsIMA2, and OsIMA1^A54V^ were fused with the N-terminal of mCherry, and driven by the CaMV 35S promoter. The combinations indicated were infiltrated into *N. benthamiana* leaves, and kept in the dark for 48 h. Then, the infiltrated leaves were used for protein extraction by RIPA buffer (50 mM Tris, 150 mM NaCl, 1% NP-40, 0.5% Sodium deoxycholate, 0.1% SDS, 1 mM PMSF, 1 x protease inhibitor cocktail [pH 8.0]) and immunoblot was conducted.

### Immunoblotting

Protein was loaded on a 12% SDS-PAGE and transferred to nitrocellulose membranes. Target proteins on the membrane were detected using immunodetection and chemiluminescence. Signals on the membrane were recorded using a chemiluminescence detection machine (Tanon-5200). The antibodies used for western blot are as follows, mouse monoclonal anti-GFP (Abmart), anti-mCherry (Abmart) and goat anti-rabbit IgG horseradish peroxidase (Affinity Biosciences).

## Acknowledgements

We thank the Institutional Center for Shared Technologies and Facilities of Xishuangbanna Tropical Botanical Garden, CAS for assistance in the determination of metal contents. We also thank Germplasm Bank of Wild Species in Southwest China for confocal laser scanning microscopy, and Crops Conservation and Breeding Base of XTBG for rice planting. This work was supported by the Applied Basic Research Project of Yunnan Province (2019FB028 and 202001AT070131).

## Author contributions

GL designed the experiments; FP, CYL, CKL and YL performed the experiments; FP, CYL, CKL, YL, PX and GL analyzed the data; FP and GL wrote the manuscript; all the authors read and approved the final manuscript.

## Supplementary data

**Supplemental Figure S1.** Identification of *OsIMA1* overexpression plants.

**Supplemental Figure S2.** Generation of *aIMA* overexpression plants.

**Supplementary Table S1**. Primers used in this paper.

