## Supplementaf information for "IRONMAN interacts with OsHRZ1 and OsHRZ2 to maintain Fe homeostasis"

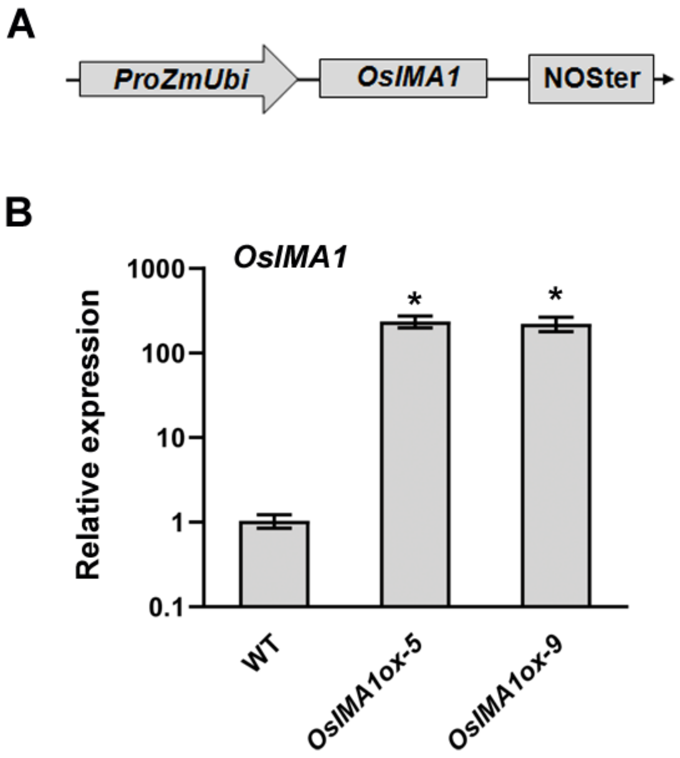


**Supplementary Figure S1.** Identification of *OsIMA1* overexpressing plants.

(A) Construction of *OsIMA1* overexpression vector. *Zea mays* ubiquitin promoter was used to drive the full-length *OsIMA1*.

(B) Expression of *OsIMA1*. Two-week-old plants were grown in Fe sufficient solution and leaves were used for qRT-PCR. Data represent means ± standard deviation (SD) ($n$ = 3). The asterisk indicates a significant difference from the wild type as determined by Student's t Test ($P < 0.05$).

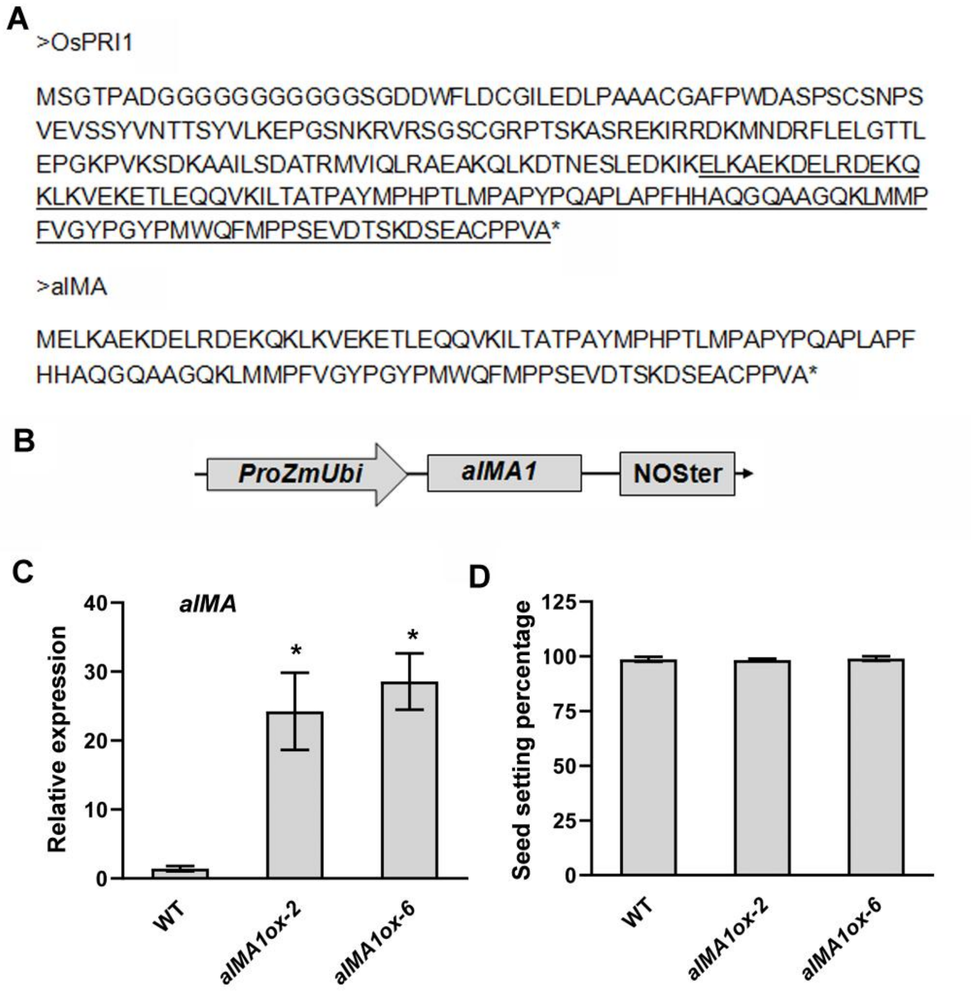


**Supplementary Figure S2.** Generation of *aIMA* overexpressing plants.

(A) Protein sequence of aIMA. The C-terminal region of OsPRI1 (the sequence underlined) was used to generate aIMA. A translation start site (M) was added.

(B) Construction of *aIMA* overexpression vector. *Zea mays* ubiquitin promoter was used to drive a*IMA*.

(C) Expression of *aIMA*. Two-week-old plants were grown in Fe sufficient solution and leaves were used for qRT-PCR. Data represent means ± standard deviation (SD) ($n$ = 3). The asterisk indicates a significant difference from the wild type as determined by Student’s t Test ($P < 0.05$).

(D) Seed setting percentage. Data represent means ± standard deviation (SD) ($n$ = 3).

**Supplementary Table S1**. Primers used in this paper.

| Name | Sequence | Vector |
|---|---|---|
| **Overexpression** | | |
| pUN-OsIMA1-F | ACTTCTGCAGGTCGACTCTAGAG ATGGCGCCGGTGTCGGAGGC | pUN1301 |
| pUN-OsIMA1-R | GAAATTCGAGCTCGGTACCCGGG TCAGGCGGCAGGCGCGCAGT | |
| pUN-aIMA1-F | ACTTCTGCAGGTCGACTCTAGAG atgGAACTGAAGGCAGAGAAGG ACG | pUN1301 |
| pUN-aIMA1-R | GAAATTCGAGCTCGGTACCCGGG TTACGCGACAGGCGGGCACGC | |
| **Yeast-two-hybrid** | | |
| yhOsHRZ1c-F | TTTcatATGTTCAAGCCTGGATGGA AGGA | GBK and GAD |
| ybOsHRZ1-R | AAAgaattCTAGTTTGGCGTAGAAC AATCTGCTG | |
| yhOsHRZ2c-F | AAAgaattcCAGAGTGAGCTTGAGG CTGAGATAC | GBK and GAD |
| yhOsHRZ2-R | TTTggatccTTAATCTGATGTTGAAC AATCAGCTCTATC | |
| yhOsIMA1-F | CTGATCTCAGAGGAGGACCTGCA tATGGCGCCGGTGTCGGAGG | GBK |
| yhOsIMA1-R | GGGAATTCGGCCTCCATGGCCAT CAGGCGGCAGGCGCGCAG | |
| yhOsIMA1-A54V-R | GGGAATTCGGCCTCCATGGCCAT CAGaCGGCAGGCGCGCAGTCATA G | GBK |
| yhOsIMA1n-R | GGGAATTCGGCCTCCATGGCCAT CAGGCCATCCTACCCCGCGCAC | GBK |
| yhOsIMA1c-F | CTGATCTCAGAGGAGGACCTGCA tGAGACAGACGGCGACGACGA | GBK |
| yhOsIMA2-F | CTGATCTCAGAGGAGGACCTGCA tATGGCGATAGCGAAGAGCGA | GBK |
| yhOsIMA2-R | GGGAATTCGGCCTCCATGGCCAT CAGGCAGCTGGAGCCACAT | |
| yhOsIMA2-A43V-R | GGGAATTCGGCCTCCATGGCCAT CAGaCAGCTGGAGCCACATCGAC G | GBK |
| yhOsIMA2n-R | GGGAATTCGGCCTCCATGGCCAT CAACCTCCTCGTCGGCAACCGT | GBK |
| yhOsIMA2c-F | CTGATCTCAGAGGAGGACCTGCA tCGCAGCAACGGCGACGACGA | GBK |

| | | |
|---|---|---|
| yhOsPRI1-F | ACGACGTACCAGATTACGCTCAtATGTCCGGTACCCCGGCGGA | GAD |
| yhOsPRI1n-R | GAATTCACTGGCCTCCATGGCCACAAGACCCTGACCTTACACGTT | |
| yhOsPRI1c-F | ACGACGTACCAGATTACGCTCAtGAACTGAAGGCAGAGAAGGACG | GAD |
| yhOsPRI1c-R | GAATTCACTGGCCTCCATGGCCATTACGCGACAGGCGGGCACGC | |
| yhOsPRI1m-F | ACGACGTACCAGATTACGCTCAtGGGTCTTGTGGTAGGCCAAC | GAD |
| yhOsPRI1m-R | GAATTCACTGGCCTCCATGGCCACAGTTCTTTAATCTTATCTTC | |
| yhOsPRI1-A253V-R | GAATTCACTGGCCTCCATGGCCATTACaCGACAGGCGGGCACGCTTCG | GAD |
| yhOsPRI4-F | AAAgaattcATGGCCTCCCCGGAGGGCTCC | GAD |
| yhOsPRI4-R | AAAggatccTTATGCAACAGGAGGGCATGACTTGG | |
| **Tripartite split-GFP** | | |
| pTG10-MYC-F | GTTGGGTCTGGCGGTGGCTCCGAGGAGCAGAAGCTGATCTCAG | pTG10-OsIMA1/2 |
| pTG10-GBK-R | GGAGGCCTGGATCGACTAGTCGGTTATGCTAGTTATGCGGCC | |
| pTG11-MYC-F | gctgaagctagtcgactctagccATGGAGGAGCAGAAGCTGATC | pTG11-OsHRZ1/2 |
| pTG11-OsHRZ1-R | actggttgatccgccaccagaccctccaccGTTTGGCGTAGAACAATCTGCT | |
| pTG11-OsHRZ2-R | actggttgatccgccaccagaccctccaccATCTGATGTTGAACAATCAGCTCTA | |
| **Degradation assays** | | |
| mCherry-OsIMA1-F | CGGCATGGACGAGCTGTACAAGATGGCGCCGGTGTCGGAGG | p30-mCherry |
| mCherry-OsIMA1-R | CATGCCTGCAGGTCGACTCTAGAGTCAGGCGGCAGGCGCGCAGT | |
| mCherry-OsIMA1-A54V-R | CATGCCTGCAGGTCGACTCTAGAGTCAGaCGGCAGGCGCGCAGTCATAG | |
| mCherry-OsIMA2-F | CGGCATGGACGAGCTGTACAAGATGGCGATAGCGAAGAGCGA | p30-mCherry |
| mCherry-OsIMA2-R | CATGCCTGCAGGTCGACTCTAGAGTCAGGCAGCTGGAGCCACAT | |
| mCherry-aIMA-F | CGGCATGGACGAGCTGTACAAGGAACTGAAGGCAGAGAAGGACG | p30-mCherry |

| | | |
|---|---|---|
| mCherry-aIMA-R | CATGCCTGCAGGTCGACTCTAGAG TTACGCGACAGGCGGGCACGC | |
| OsHRZ1-GFP-F | ctgatcagcgaggaggacctggATGGCGAC GCCGACGCCCATG | p30-MYC-GFP |
| OsHRZ1-GFP-R | cttgctcaccattctagaggtgTTTGGCGTAG AACAATCTGCTGT | |
| OsHRZ2-GFP-F | ctgatcagcgaggaggacctggATGGAGAA AATCGAGAGGCAC | p30-MYC-GFP |
| OsHRZ2-GFP-R | cttgctcaccattctagaggtgTCTGATGTTG AACAATCAGCT | |
| **qRT-PCR** | | |
| q-OsIMA1-F | CACCAAGACGGCGGCATTAT | |
| q-OsIMA1-R | TCTGTCTCGGCCATCCTACC | |
| q-OsIRO3-F | GCGAGCTGGGTAATATGCTAGA | |
| q-OsIRO3-R | ATCCGGGTGGTGTCAGTTAG | |
| q-OsIRO2-F | GAAGGTCTTCACTTCATCAGTTCA | |
| q-OsIRO2-R | TGATCGTTCCTTCACTTCTCTG | |
| q-OsNAS1-F | GTGGTTCTGCCGGTGGTC | |
| q-OsNAS1-R | AGACGGACAGCTCCTTGTTG | |
| q-OsNAS2-F | CGTCTGAGTGCGTGCATAGT | |
| q-OsNAS2-R | CACAAACACAAACCGATACCA | |
| q-OsYSL15-F | ATCTCACCTTGACATCGCCG | |
| q-OsYSL15-R | AAACGCCCTGTAGAACAGCA | |
| q-OsTOM1-F | AGAGGTGCTGCAAATGGCATAT | |
| q-OsTOM1-R | ATCATTTGATCCCCTGGGAAGA | |
| q-OsACTIN1-F | ACACCGGTGTCATGGTCGG | |
| q-OsACTIN1-R | ACACGGAGCTCGTTGTAGAA | |
| q-OsOBP-F | GGCGCCGAAGAAGCTATTG | |
| q-OsOBP-R | GTTGCTCTTCAAGCCTGCTC | |
